# Improving Drug Sensitivity Prediction and Inference by Multi-Task Learning

**DOI:** 10.1101/2024.05.09.593186

**Authors:** Jared Strauch, Amir Asiaee

## Abstract

The development of models to predict sensitivity to anticancer drugs is an area of significant interest, given the diverse responses to treatment among patients and the considerable expense and time involved in anticancer drug development. Leveraging “omic” data and anticancer response information from the Cancer Cell Line Encyclopedia, we propose a novel approach utilizing multitask learning to enhance prediction accuracy and inference. We extended a multitask learning framework called the Data Shared Lasso to develop the Data Shared Elastic Net. This enabled the construction of tissue-specific models with information sharing while maintaining the attractive properties of Elastic Net regression. By employing this approach, we observed improvements in prediction accuracy compared to single-task Elastic Net models, particularly for cell lines displaying high sensitivity to treatment. Furthermore, the Data Shared Elastic Net facilitated the identification of predictors for anticancer drug sensitivity within specific tissue types, shedding light on cellular pathways targeted by these drugs across tissues. We also investigated the impact of data leakage on modeling outcomes from previous studies, which led to underestimating prediction error and erroneous inferences

## Introduction

The development of models to predict sensitivity to anticancer drugs is an area of significant interest, given the diverse responses to treatment among patients and the considerable expense and time involved in anticancer drug development. Accurately predicting drug sensitivity has the potential to improve patient care dramatically, reduce the time and cost of developing new treatments, and even offer opportunities for repurposing already approved anticancer drugs. Several comprehensive public datasets, such as the Cancer Cell Line Encyclopedia (CCLE), Genomics of Drug Cancer Sensitivity (GDSC), and Cancer Therapeutics Response Portal (CTRPv2), have been created to foster the development of such predictive models. These datasets contain various “omic” measurements alongside drug sensitivity measurements.

Our work primarily revolves around the analysis of CCLE data, although the methods used could be applied to other large “omic” datasets. However, a primary challenge in developing accurate prediction models using these datasets is the scarcity of cell lines demonstrating sensitivity to individual drugs—due to anticancer drugs typically being specialized for specific types of cancer rather than broad-spectrum. Consequently, models may be overly conservative in predicting sensitivity for specific cell lines, given the preponderance of insensitive responses to the chosen treatment. To address this issue, we propose a multitask learning framework, where we view modeling within each tissue type as individual tasks and leverage shared information between these tasks.

### Dataset and Previous Work

The Cancer Cell Line Encyclopedia (CCLE) is a comprehensive genomic dataset that was initially released in 2012. It comprises genomic information for 947 human cancer cell lines and the pharmacological profiles of 24 compounds tested across approximately 500 of these lines. The CCLE covers 36 different types of tumors. Each cell line has been extensively characterized using various genomic technology platforms.

The mutational status of over 1,600 genes was determined through targeted massively parallel sequencing, with subsequent removal of variants likely to be germline events. Additionally, 392 recurrent mutations affecting 33 known cancer genes were evaluated by mass spectrometric genotyping. DNA copy number was measured using high-density single nucleotide polymorphism arrays (Affymetrix SNP 6.0), and messenger RNA expression levels were obtained for each line using Affymetrix U133 plus 2.0 arrays.

Elastic Net regression modeling demonstrated that known drug targets could be identified as significant predictors of drug sensitivity. Furthermore, previously unidentified predictors were also discovered. For instance, plasma cell lineage correlated with sensitivity to IGF1 receptor inhibitors; AHR expression was associated with MEK inhibitor efficacy in NRAS-mutant lines; and SLFN11 expression predicted sensitivity to topoisomerase inhibitors.

In 2019, the CCLE was expanded to include genetic, RNA splicing, DNA methylation, histone H3 modification, microRNA expression, and reverse-phase protein array data for 1,072 cell lines from individuals of various lineages and ethnicities. Drug response information was obtained from the GDSC database, which includes 265 anticancer drugs and two dose-response measures: Inhibitory Concentration 50 (IC50) and Area Under the Curve (AUC), Figure 1. The AUC was selected as the response of interest due to its robustness to outliers, while the IC50, a point estimate from a fitted curve, is more susceptible to influence. Adding reverse-phase protein array (RPPA) measurements as features enhanced the prediction performance of the Elastic Net regression model as measured by the Mean Squared Error (MSE).

**Fig. 1.**
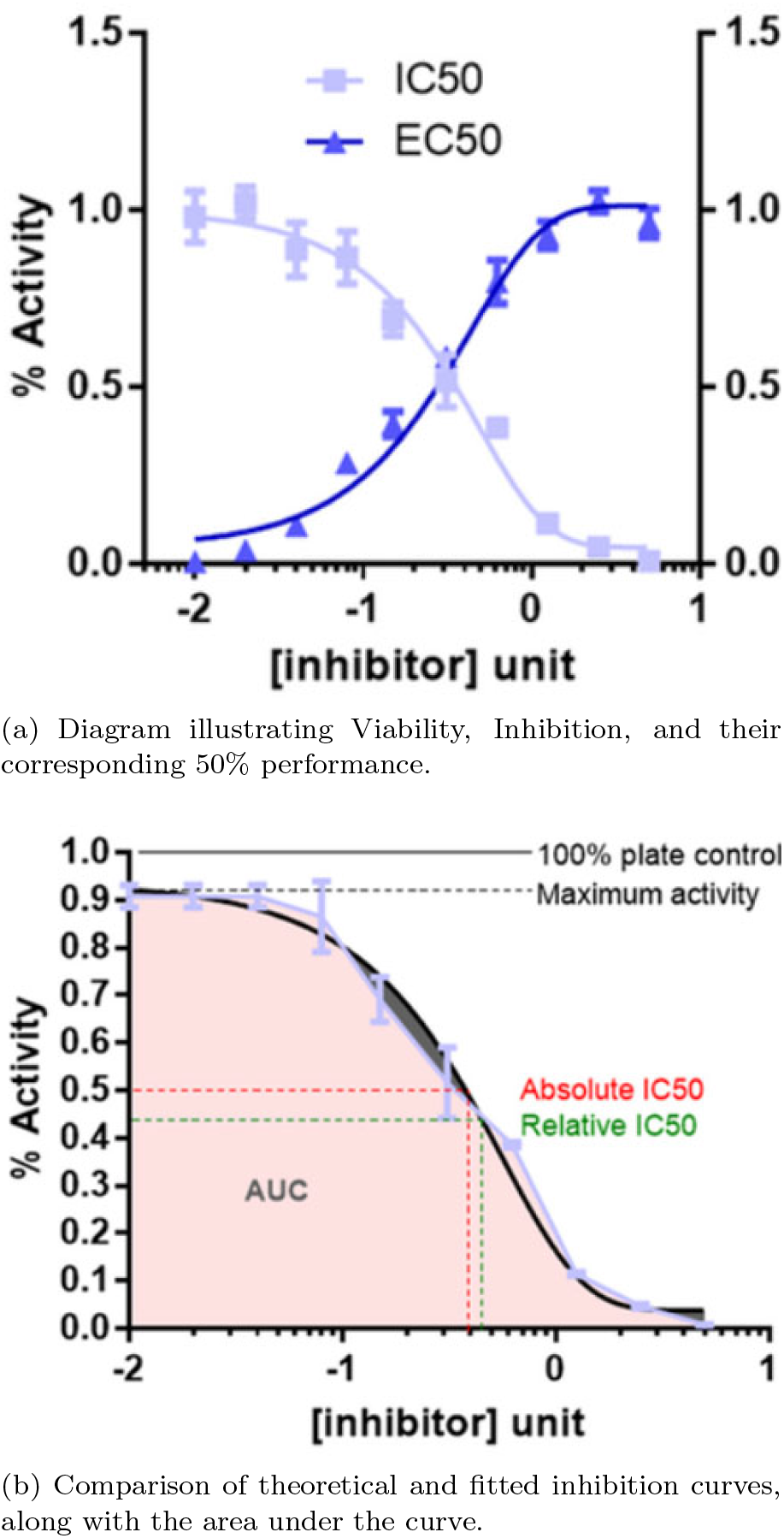
Definition of IC50, EC50, and AUC from dose-response plots [20]. (a) Dose-response plots showcasing EC50 (effective concentration required to achieve 50% activity - depicted in dark blue) and IC50 (inhibitor concentration leading to 50% inhibition - shown in light blue). (b) Estimation of the area under the inhibition curve (AUC), with the theoretical (red) and fitted (green) IC50 measurements from the dose-response curves. The empirical light blue curve is typically fitted using a parametric model (such as the Hill equation), with a maximum of three biological replicates per concentration. The theoretical activity (solid black line) is represented as plate control, and the maximum activity tested (broken black line) can vary.

### Modeling

#### LASSO Regression

LASSO (Least Absolute Shrinkage and Selection Operator) regression, introduced by [16], is a linear regression technique that incorporates shrinkage. Shrinkage involves reducing the magnitude of certain regression coefficients toward zero. In LASSO regression, shrinkage is achieved through an *l*_1_-norm penalty, resulting in both regularization and feature selection. The *l*_1_-norm penalty promotes simpler and sparser models by encouraging coefficients to be set exactly to zero. This characteristic makes LASSO regression particularly valuable for variable selection in situations where the number of predictors (*p*) is much larger than the number of observations (*n*), such as high-dimensional datasets. The objective function to be minimized in LASSO regression can be represented as follows:

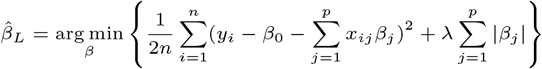

In this formula, the first term is the residual sum of squares, and the second term is the sum of the absolute values of the model parameters, which is weighted by the regularization parameter, *λ*. The regularization parameter controls the amount of shrinkage: the larger the value of *λ*, the greater the shrinkage.

#### Elastic Net Regression

Elastic Net (EN) regression, [21], is a regularization method that linearly combines the *l*_1_ and *l*_2_-norm penalties of the Lasso and Ridge methods. Like Lasso, it can reduce some coefficients exactly to zero, performing variable selection. However, Elastic Net overcomes one limitation of the Lasso: its tendency to select only one variable from a group of highly correlated variables. By incorporating the Ridge (*l*_2_ norm) penalty [11], Elastic Net can include all correlated variables in the model, making it particularly useful when dealing with multicollinearity. This is desirable in genomics settings where genes in a pathway tend to be correlated, and the whole pathway is predictive of the outcome. In these circumstances, we would like to select all genes in the pathway in the final model instead of only one gene representing the pathway.

The objective function for Elastic Net regression is a combination of the objective functions of Lasso and Ridge, and it can be represented as follows:

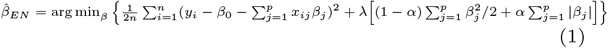

In this formula, the first term is the residual sum of squares, the second is the Ridge penalty (*l*_2_ norm), and the third is the Lasso penalty (*l*_1_ norm). The parameter *λ* controls the overall strength of the penalty, and *α* determines the mixing proportion of *l*_1_ and *l*_2_ penalties. When *α* = 1, Elastic Net is equivalent to Lasso, and when *α* = 0, it is equivalent to Ridge.

#### Multitask Learning

Multi-task learning (MTL) is a subfield of machine learning where multiple learning tasks are solved simultaneously while exploiting commonalities and differences across tasks. This is in contrast to traditional machine learning approaches that handle tasks independently. The key idea behind MTL is that by learning tasks in parallel, the algorithm can leverage the information contained in multiple related tasks to improve the generalization performance. This is particularly beneficial when the tasks are related in some way, as the learning algorithm can transfer knowledge from tasks with more data to tasks with less data.

Mathematically, multi-task learning can be formulated as an optimization problem. Suppose we have *T* tasks, and for each task, *t*, we have a loss function *L*_*t*_(*β*_*t*_, *X*_*t*_, *Y*_*t*_), where *β*_*t*_ is the parameter of interest to be learned, *X*_*t*_ is the input data, and *Y*_*t*_ is the output data for task *t*. The goal is to minimize the total loss across all tasks, possibly subject to some constraints that encourage the tasks to share information. This can be written as:

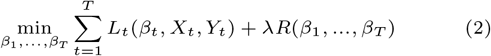

In this formula, *R*(*β*_1_, …, *β*_*T*_) is a regularization term that encourages the vectors *β*_1_, …, *β*_*T*_ to be similar in some sense, and *λ* is a hyperparameter that controls the strength of the regularization. The specific form of the regularization term *R* depends on our assumptions about the tasks. For example, assuming all tasks share a common feature representation, we might choose *R* to encourage the coefficient vectors to be close in Euclidean distance.

In the realm of drug sensitivity prediction, we consider each task as the prediction of drug response for a specific tissue, denoted as *t*. This approach is motivated by the observation that lineage (the tissue from which the cell line is derived in the CCLE dataset) is a significant predictor [3]. Rather than using lineage as a predictor, we propose to create a distinct regression model for each tissue. This approach offers more flexibility than a single regression model for all cell lines. However, there might be a universal omics feature signature that aids in predicting drug sensitivity, making it beneficial to have a flexible model that allows for coefficient sharing between tissues. We suggest the use of Data Shared Lasso/Elastic Net regression within a multi-task learning framework to facilitate information sharing between tissue types and strike a balance between enhancing flexibility to improve prediction accuracy and maintaining interpretability, as discussed in Section 2.2.

It is essential to clarify how our method relates to other multi-task learning frameworks specifically designed for drug response prediction. The primary concern lies in the use of the term “multi-task learning,” which is often used instead of “multi-output prediction” in most related work. In multi-output prediction, a single feature matrix is used to predict multiple outcomes, and the relation between the outcomes can enhance overall prediction performance. In drug response prediction, each output is an AUC of a drug, and a single feature matrix of all drugs serves as the predictor matrix. One parameter is learned per outcome as:

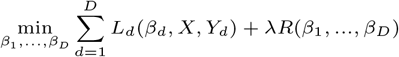

In this equation, the regularizer function *R* links the *D* prediction problems together and aids in information borrowing. It’s important to note that here, the loss function *L*_*d*_ is a function of all samples *X*, while in multi-task learning (2), the loss is a function of task-specific samples *X*_*t*_. Therefore, many papers have mislabeled their proposed multi-output learning model as multi-task learning. The methods of information sharing between drugs vary, and it’s beyond the scope of this report to cover all methods in detail. Kernelized Bayesian multitask learning [4] and Trace Norm regularization [19] are two examples of linear regression-based multitask learning used to share information between models fit for anticancer drugs. Both methods have improved prediction accuracy compared to Elastic Net regression per drug.

While there are many instances of multitask/multi-output learning being used to share information between drugs, we have not found any articles that explicitly treat each tissue type as a task for information sharing. This does not imply that other modeling strategies have not utilized tissue information. [12] developed the Tissue-guided LASSO, which trains a LASSO regression model on all cell lines not originating from the tissue of interest and selects the hyperparameter of the LASSO model, *λ*, that results in the best performance in the tissue of interest. In other words, instead of using K-fold cross-validation to tune the hyperparameter *λ*, they use tissue labels to split the data into the tissue of interest that the model is being developed for (test) and the rest of the samples from other tissues (train). They demonstrate that a model trained this way using CCLE performs better on unobserved clinical outcome data than other methods.

Numerous other multitask learning methods indirectly incorporate tissue information by determining cell line similarity through pairwise correlation of cell line features, typically restricted to gene expression measurements. Essentially, these methods cluster cell lines based on their gene expression and treat drug sensitivity prediction in each cluster as a separate task. This approach indirectly employs tissue type information because cell lines from the same tissue tend to exhibit greater similarity to each other than to cell lines from different tissues. These methods are comprehensively reviewed in [5]. While these methods offer substantial flexibility and significantly enhance prediction accuracy, the amalgamation of clustering and prediction can complicate the interpretation of results. We show subsequently that Data Shared Elastic Net regression achieves a balance between flexibility and interpretability, enabling improved prediction accuracy without sacrificing ease of interpretation.

## Methods

In this section, we first outline our efforts to reproduce the findings of the original CCLE papers and discuss the issues we encountered with their implementation. Subsequently, we introduce our multi-task learning framework.

### Replication of CCLE Results

We endeavored to reproduce the results from [7], utilizing mutation, gene copy number, RNA expression, and RPPA data to construct our feature matrix *X* and AUC as our response vector *y*. The general pipeline they employed is as follows:

- **Data preparation:**
  - Compile the feature matrix and response vector,
  - Limit the data to cell lines with observed responses and at least one observed predictor
  - Substitute missing predictor values with zeroes,
  - Perform feature filtering (See Section 2.1.1)
- - **Model fitting:**
  - Apply 10-fold cross-validation to tune the parameters for the Elastic Net regression model (both *α* and *λ*)
  - Identify the optimal parameter set, i.e., (*α*^*\**^, *λ*^*\**^) that yields the best performance on the test set.
- - **Inference:**
  - Refit the model 200 times with the tuned parameters (*α*^*\**^, *λ*^*\**^) using 200 bootstrapped datasets learning 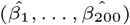 coefficient vectors.
  - From the bootstrapped models, average the estimated coefficients to ascertain the direction and magnitude of the effect on the predicted sensitivity: 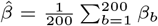
  - Use the proportion of times each feature’s estimated coefficient was non-zero as a general measure of the feature’s importance (score) in predicting drug sensitivity: 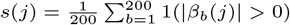

We obtained the data from the CCLE web portal and adhered to the data processing steps outlined in the methods section from [3] and [7]. However, we were unable to reproduce Figure 11-i of [7] from raw data. Upon contacting M. Ghandi, they provided us with the code and data used to generate the figure. Upon reviewing their code, we discovered that their data processing led to data leakage during cross-validation, which explained our inability to reproduce their findings.

#### Data Leakage During Cross-validation

The feature matrix comprises approximately 80,000 predictors, with typically between 200-600 observations for each anticancer drug. The authors of the CCLE papers implemented some feature selection to reduce the size of the feature matrix, starting by eliminating all features with a variance less than 0.01. Subsequently, they calculated pairwise correlations between each feature and the response vector, removing all features with a correlation less than 0.1. Features were also standardized to ensure that no feature was penalized more or less heavily due to the measurement scale. However, all of these processing steps were performed outside of the 10-fold cross-validation used to tune the Elastic Net regression model parameters, leading to information leakage between the test and train folds. This resulted in an underestimation of the prediction error during cross-validation and overfitting in the model.

To rectify the data leakage, we performed the same feature selection using only the training folds and then restricted the features of the testing folds to match the training folds. We also centered and scaled the testing data using the mean and standard deviation from the training data. In [7], Ponatinib is the only drug with a figure that we can use to validate the results of our replication attempt. Therefore, Ponatinib will be included as an example in all of our main results. Using their original code and data, we could perform the analysis for all of the drugs. Hence, we will also include examples from other drugs when they better highlight an important result than Ponatinib alone.

#### Consequence of Data Leakage for Prediction

Figure 2 displays the cross-validation error during parameter tuning for Ponatinib. Although data leakage does not significantly impact the cross-validation error, we observe that the values (*α*^*\**^, *λ*^*\**^) selected during parameter tuning differ greatly. This leads to substantial differences in the degree of coefficient shrinkage in each model and dramatically affects the results. Figure 3 shows the cross-validation error for Bleomycin. This figure demonstrates that while the selected parameters are very similar, the cross-validation error for these parameters varies significantly. Collectively, the two examples show that data leakage is not guaranteed to lead to underestimating cross-validation errors or selecting different values during parameter tuning, but we can see that it does occur.

**Fig. 2.**
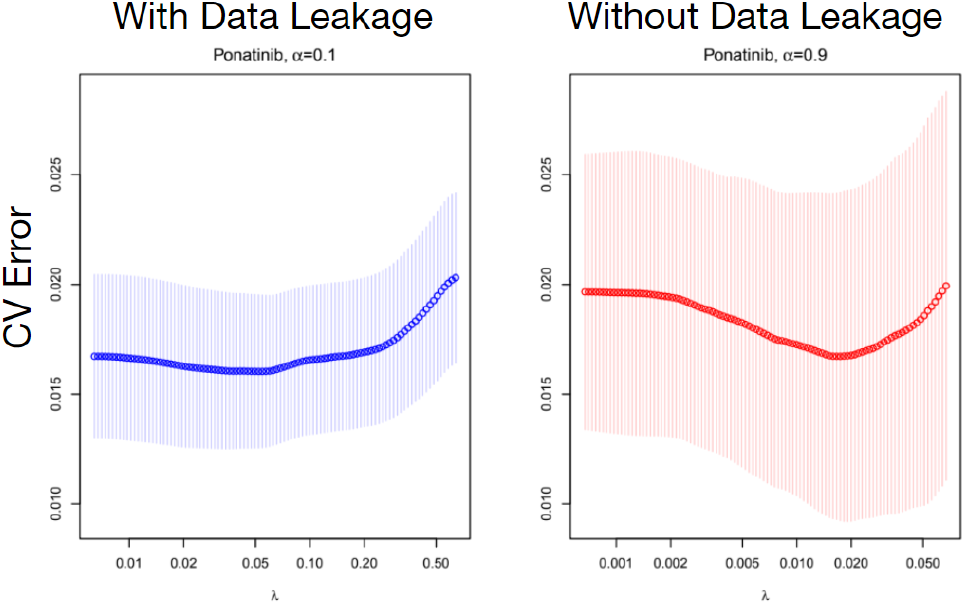
Comparing Cross-Validation Error for Ponatinib. While the optimal cross-validation errors are the same, the tuned parameters are not: (*α*^*\**^ = 0.1, *λ*^*\**^ = 0.055)*≠* (*α*^*\**^ = 0.9, *λ*^*\**^ = 0.020)

**Fig. 3.**
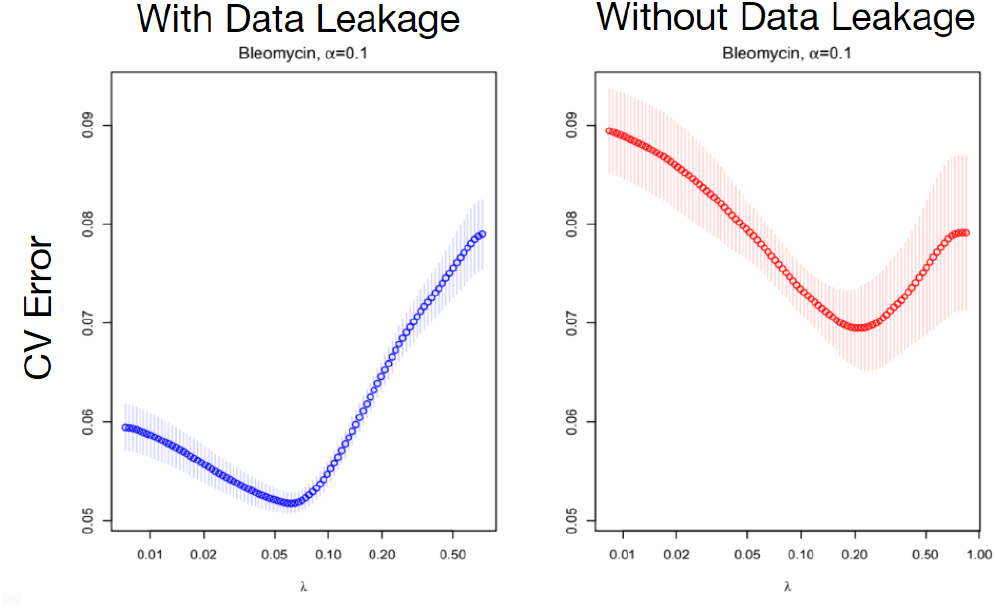
Comparing Cross-Validation Error for Bleomycin. Here the optimal cross-validation errors (minimum of the CV curves) are very different.

#### Consequence of Data Leakage for Inference

To measure feature importance, [3] and [7] used the mean coefficient estimate 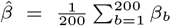 and the proportion of times each feature was non-zero in bootstrap replicates of the model 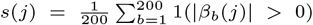, Figure 4. A feature *j* is important and predictive according to [3] if its score *s*(*j*) *>* 0.8, i.e, it shows up as non-zero in 80% runs for Elastic Net on 200 bootstrapped datasets. Figure 4 demonstrates the difference in features and mean coefficient estimates that meet Ponatinib’s 0.8 non-zero proportion threshold with and without data leakage. We see that only one feature meets this threshold for the model without data leakage and six for the model with data leakage. Though both models agree on SHP-2 expression being the most important feature, the estimated coefficient differs in each model.

**Fig. 4.**
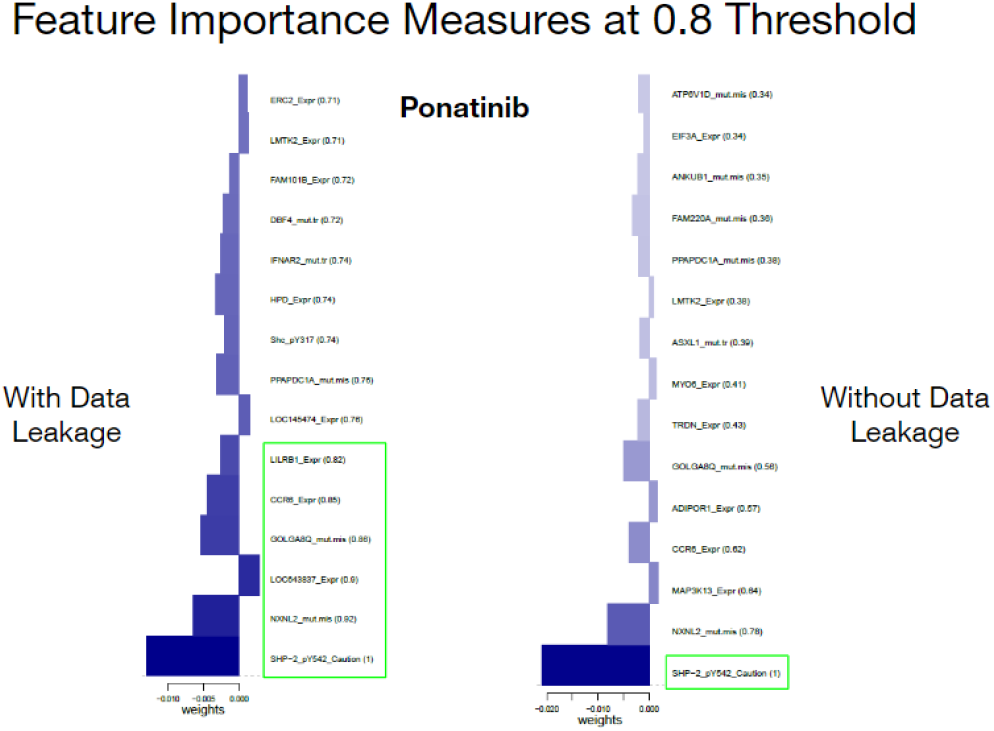
Comparison of Feature Importance for Sensitivity to Ponatinib. The numbers in parentheses in front of each feature represent the proportion of times that the feature had a non-zero coefficient in Elastic Net estimation for 200 bootstrapped datasets. The green rectangles highlight features surpassing the 0.8 importance threshold, while the bars indicate the coefficient values.

It is important to note that the similarity of the important feature list between the scenarios with and without data leakage, as indicated by the green rectangles in Figure 4, is dependent on the chosen importance threshold. To see the discrepancy between the methods as a function of threshold, we used the Jaccard index. Given two lists, *A* and *B*, the Jaccard index computes their similarity as 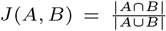 and therefore 0 ≤ *J*(*A, B*) ≤ 1. Using the threshold that generated the two lists as a superscript, we have 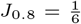 for Figure 4. We computed the Jaccard index for *i* = [100]*/*100 threshold values and plotted *J*_*i*_, along with the number of features above the threshold for each method. A Jaccard Index of zero indicates complete disagreement. As the Jaccard index increases to one, there is more agreement between the two methods. Figure 5 shows that for Ponatinib, there is little agreement between the methods until a threshold value of 0.9 when there is only one feature for the model without data leakage and two features for the model with data leakage. For many of the threshold values, there is around an order of magnitude difference between the number of features meeting the threshold for each model, i.e., the Elastic Net model with data leakage incorrectly detects more predictive features.

**Fig. 5.**
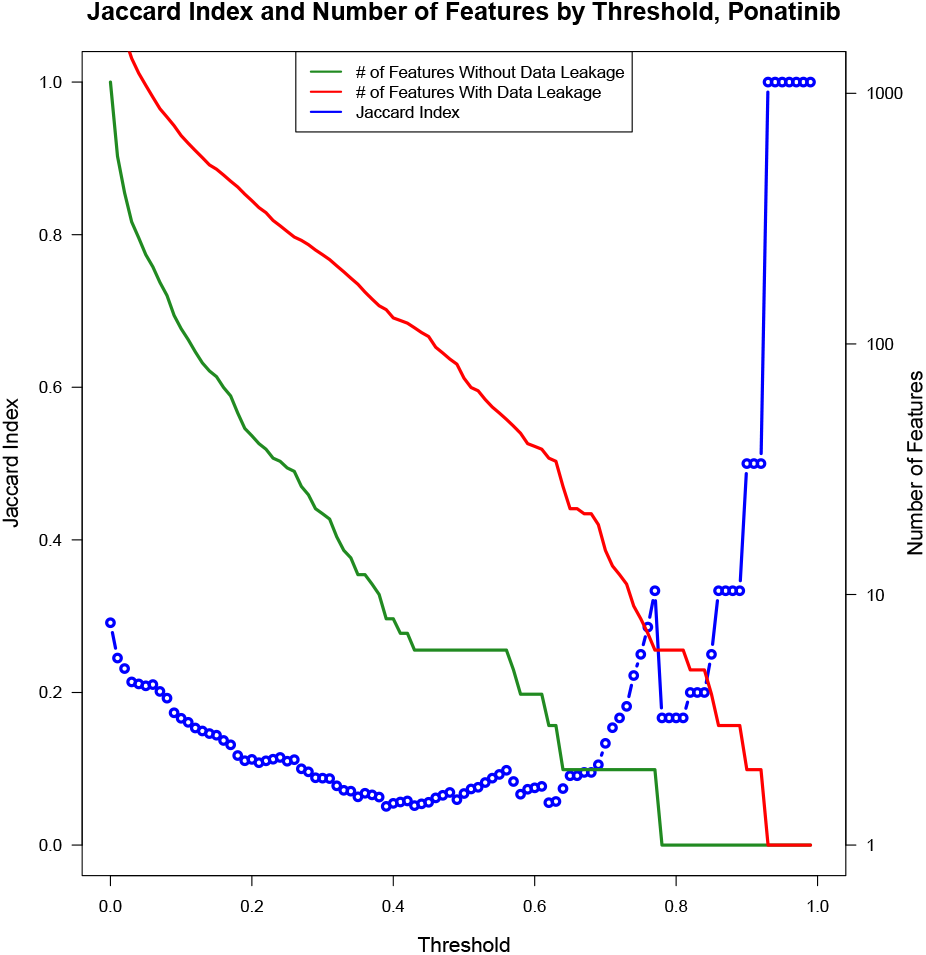
Jaccard Similarity of Predictive Features Selected with and without Data Leakage for Ponatinib at Various Non-zero Importance Thresholds. The number of features selected by each method at a threshold is presented on a log_10_ scale, right-hand side. There is a large amount of disagreement between the methods for thresholds *<* 0.9.

To summarize how data leakage affects our assessment of feature importance for all drugs, we computed the Jaccard index for each drug with a threshold of 0.8 and plotted a histogram of the values in Figure 6. The mean Jaccard index for all drugs with a non-zero proportion threshold of 0.8 was 0.180 with a standard deviation of 0.302, indicating little agreement between the two methods on average. This disagreement between the methods can lead to very different inferential results and highlights the importance of ensuring there is no data leakage when performing cross-validation.

**Fig. 6.**
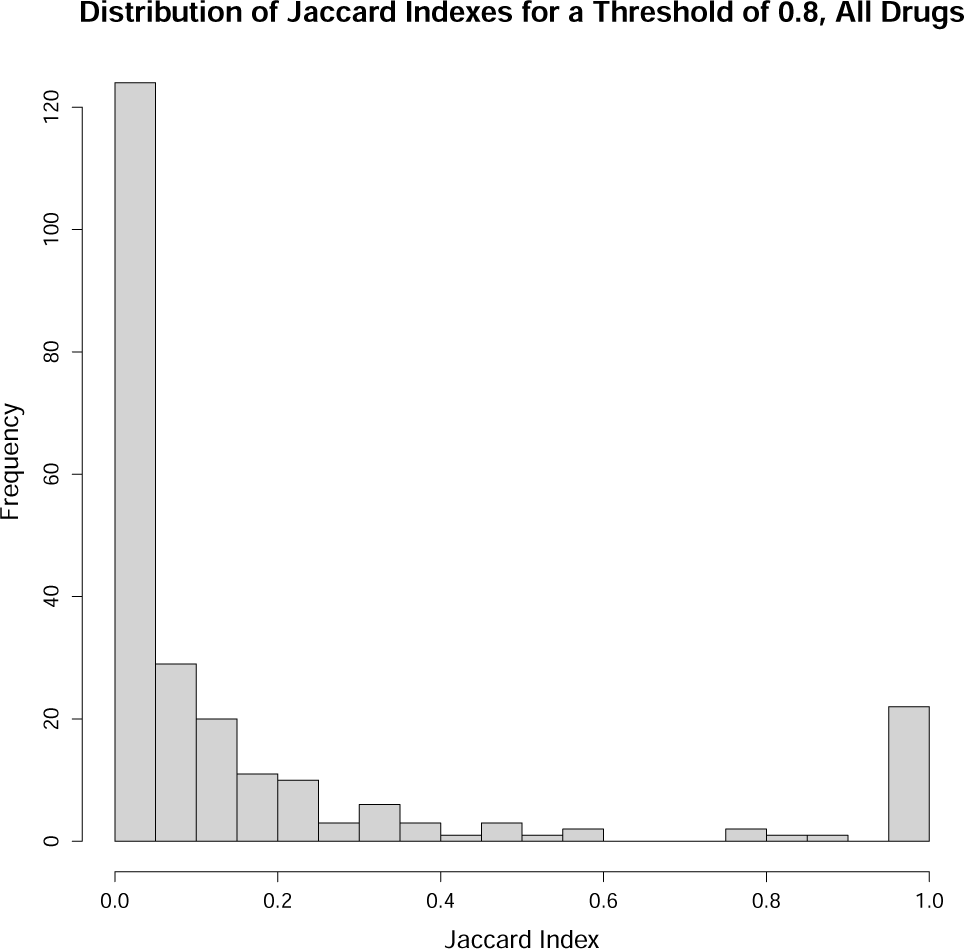
Jaccard Similarity between Important Feature Lists Computed with and without Data Leakage for All Drugs With 0.8 Importance Threshold. We see that a majority of drugs disagree on which features are important at this threshold.

### Proposed Methods

After correcting the data leakage in the modeling strategy of [7], we explored methods to enhance the prediction accuracy and inference generated compared to Elastic Net regression. In [3], the tissue types of the cell lines were incorporated as one hot-coded (a binary vector) features. Tissue lineage was identified as a significant predictor for the drugs AEW451, AZD0530, LBW242, Nilotinib, PF2341066, and ZD-6474. This is biologically plausible as anticancer drugs are typically effective at treating only specific cancer types and are not universally active. Therefore, for each drug, tissue types would be expected to provide valuable information on drug sensitivity. We hypothesized that separate modeling of all cell lines originating from each tissue type might enhance prediction accuracy and enable us to identify features crucial for predicting drug sensitivity within each tissue.

One challenge with predictive modeling per individual tissue is the small number of samples within each tissue. For instance, each drug in the updated 2019 CCLE data has response information for 200-600 cell lines, with the cell lines distributed across 19 different tissue types. This means that, on average, we have 10-30 samples per tissue type. With approximately 80,000 features, this would make model fitting extremely challenging. To circumvent this issue, we employed a multi-task learning framework that shares information between tasks through a shared parameter, namely Data Shared Lasso (DSL) regression [8]. In this context, each task is defined as fitting a penalized model for a given tissue type, while a shared parameter is fitted to the entire data.

The Data Shared Lasso objective is as follows:

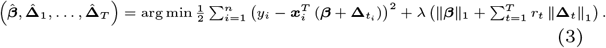

Here, *λ* is the regularization hyperparameter controlling the complexity of the parameter. The first part of the DSL objective is the Residual Sum of Squares, and the second part is the norm that induces the parameters’ sparsity. *β* is the shared parameter between tasks, and Δ_*t*_ is the corresponding parameter of task *t*. The concept is that, given sufficient data from each group, we could accurately estimate ***β*** + **Δ***t*_*i*_ as one parameter for each task *t* using separate regression procedures. Alternatively, if we lacked sufficient data, we could pool all of it together and estimate a single coefficient vector that performs best when averaged across the groups. DSL uses regularization parameters *r*_*t*_ to control the amount of pooling done in an intermediate problem.

#### Fitting the Data Shared Lasso

By rewriting the DSL objective, we can use Lasso solvers to fit it: Let *Z* be defined as

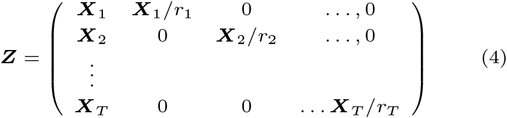

where *X*_*j*_ and *y*_*j*_ are the dataset for tissue *j*. Namely, *X*_*j*_ is the matrix formed by taking as rows all *x*_*i*_ such that *t*_*i*_ = *j* and *y*_*j*_ is the vector formed by taking as elements all *y*_*i*_ such that *t*_*i*_ = *j*. Finally, let 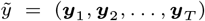and 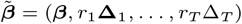 and all *r*_*t*_ *>* 0.

Then we note that

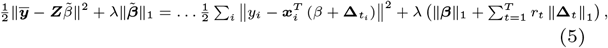

Where *β* is a shared parameter to be estimated for all tasks, and Δ_*t*_ is the deviation from the shared parameter for each task. *λ* is the classic LASSO *l*_1_ norm regularization penalty where larger values of *λ* increase shrinkage. From ***Z***, it is easy to see how *r*_*t*_ affects the amount of information shared between tasks. As *r*_*t*_ approaches zero, it is equivalent to fitting separate LASSO models for each task, and as *r*_*t*_ increases to infinity, it is equivalent to fitting one pooled LASSO model. In general, increasing the value of *r*_*t*_ increases the amount of information shared between each task. [8] give recommendations for values of *r*_*t*_ depending on which coefficient estimates are desired and are summarized in Table 1. [8] also note that if the primary interest is minimizing prediction error, then *r*_*t*_ can be tuned alongside *λ* using cross-validation. An advantage of using the LASSO objective function is that special case solvers have been developed already, which leads to more time-efficient solutions and methods that work with sparse data storage. This is important because the feature matrix requires large amounts of memory when there are many tasks.

**Table 1.** How *r*_*t*_ changes the estimate of 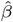.

| $r_t$ Value | Effect on $\hat{\beta}$ |
| --- | --- |
| $r_t$ such that $\sum_t r_t < 1$ | We are guaranteed that $\hat{\beta} = 0$. This is equivalent to fitting separate lasso models to each group. |
| $r_t$ such that $\sum_t r_t = 1$ | There will be identifiability issues, so this case should be avoided. |
| $r_t = r > 1 \forall t \in \{1, 2, \dots, T\}$ | $\hat{\beta}$ will be the median of $\beta + \Delta_t$. |
| $r_t = r \in \left(\frac{1}{T}, \frac{1}{T-2}\right) \forall t \in \{1, 2, \dots, T\}$ | We would have nonzero $\hat{\beta}$ if and only if all of the $\beta + \Delta_t$ are nonzero with the same sign. Then, we would have $\hat{\beta} = \text{sign}(\beta + \Delta_1) \min(\beta + \Delta_t)$. |

#### Data Shared Elastic Net

In order to make a more direct comparison on how incorporating tissue information improves prediction and inference compared to Elastic Net regression, we extended the Data Shared Lasso to include an *l*_2_ norm penalty term in the same manner as Elastic Net Regression, resulting in the objective function below.

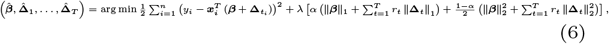

The objective is equivalent to the following:

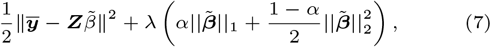

where the matrix *Z* and vector 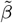 are defined as before. This reformulation lets us use the Elastic Net solver to optimize the DS Elastic Net (DSEN) objective. Theoretical properties of data sharing with general norms have been investigated in [1, 2] and recently [13] showed that DSEN regression is helpful in the integration of multiple gene expression data sets.

To implement the DSEN, we used the same dataset and pre-processing as [7] except for the feature variation and pairwise feature-response correlation cutoffs. Each task needs to have the same features to estimate the shared coefficient between each task; therefore, we cannot select features by using within-feature variation and pairwise feature-response correlation cutoffs within each tissue feature matrix. Performing the feature variation and pairwise correlation cutoff on the pooled feature matrix is unlikely to capture the true variation within features and the correlation between features and response for each task, so we omitted these steps altogether. The data was standardized after forming the Z feature matrix and separating the data into train and test folds.

Due to the complexity that would result from trying to tune *r*_*t*_, *λ*, and *α*, we elected to set *r*_*t*_ to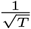 as recommended by [8] and tuned *λ* and *α* using 10-fold cross-validation. After parameter tuning was finished, we refit the model on all of the data to get estimated drug response values which were then compared to the results of Elastic Net regression through the root mean squared error (RMSE) of predicted AUC values. We also performed 200 bootstrap re-samples of the data and refit the model on each re-sample in the same manner as described in [3]. From these bootstrap replicates, we generated heatmap plots for the mean shared and individual task coefficient estimates of the 15 features with the highest proportion of non-zero shared or tissue-specific coefficient estimates in the refit models.

## Results

### Comparing Prediction Accuracy Between DSEN and EN Regression

The evaluation of model performance revealed that the DSEN model exhibited a lower root mean square error (RMSE) compared to the EN model for 13 out of the 16 drugs tested, as illustrated in Figure 7.

**Fig. 7.**
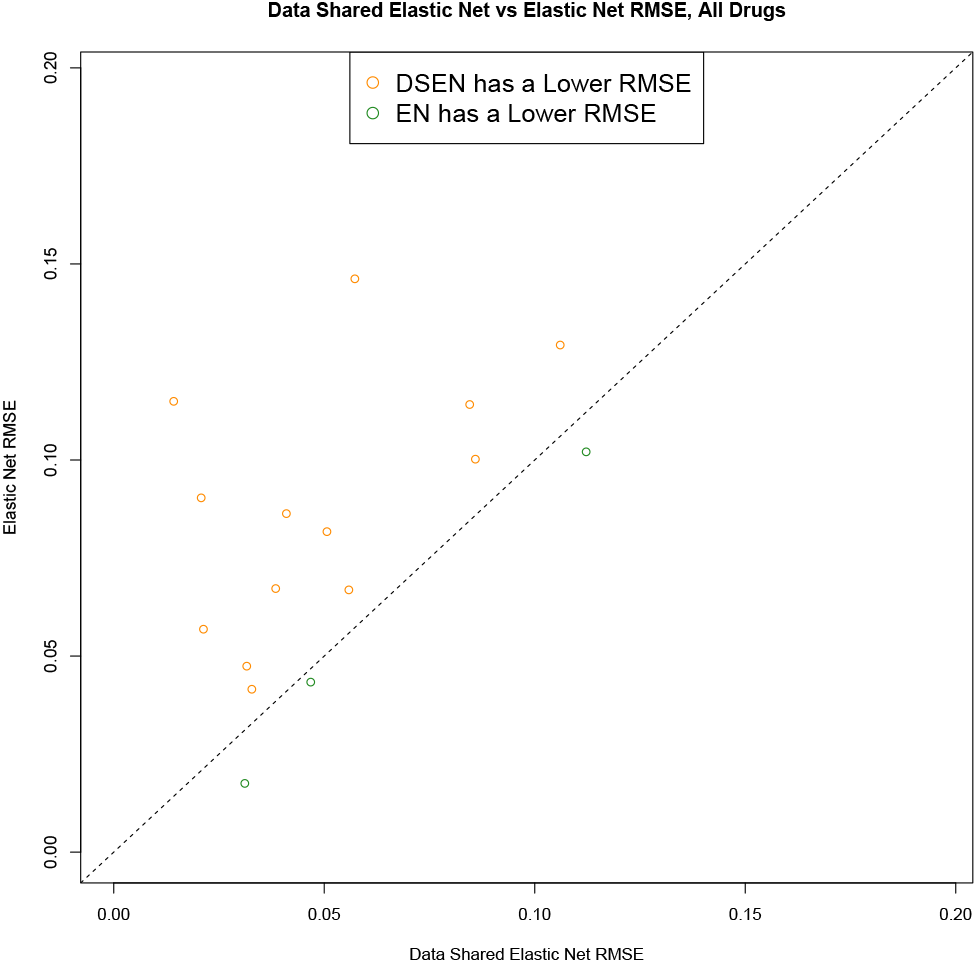
Comparing RMSE for Predicted AUC values from the DSEN and EN models for All Drugs. Points above the y = x line indicate the DSEN model has a lower RMSE. Points below the y = x line indicate the Elastic Net model has a lower RMSE. The DSEN model outperforms the EN model for nearly every drug.

Specifically, DSEN regression outperformed EN regression in prediction accuracy for tissues sensitive to the targeted anticancer drug. A prominent example is Ponatinib, which is used to treat adult chronic myeloid leukemia or Philadelphia chromosome-positive acute lymphoblastic leukemia that has shown resistance or intolerance to prior tyrosine kinase inhibitor therapy. When comparing the RMSE values for cell lines derived from different tissue lineages, the most substantial improvement between the DSEN and EN models was observed in leukemic cell lines, as depicted in Figure 8. This pattern was consistent for other drugs. In PLX-4720 and Tanespimycin, two drugs that target melanoma, the largest difference in RMSE values were observed in cell lines derived from skin tissues.

**Fig. 8.**
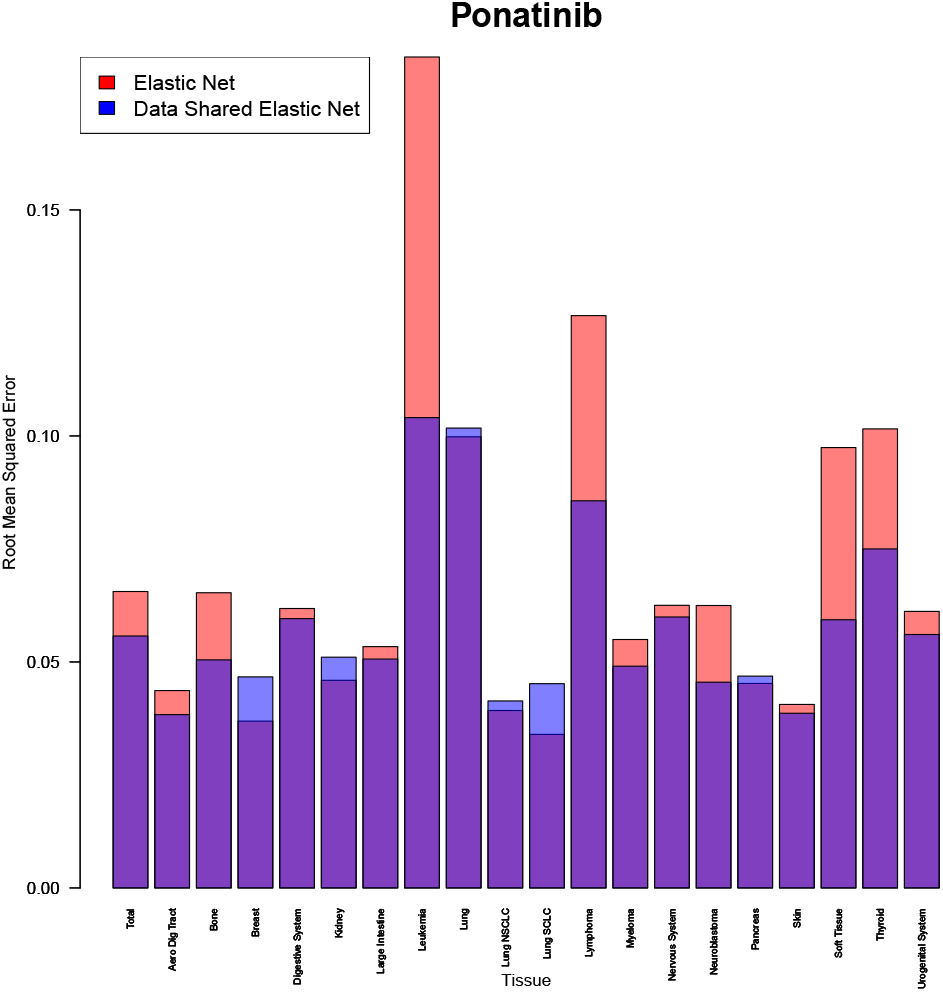
Comparison of RMSE between the DSEN and EN models for Predicted AUC values of Ponatinib. RMSE values were computed for all cell lines as well as within specific tissue types. Notably, the highest difference in RMSE values was observed in cell lines derived from leukemic tissues, representing the primary target cancer of Ponatinib.

**Fig. 9.**
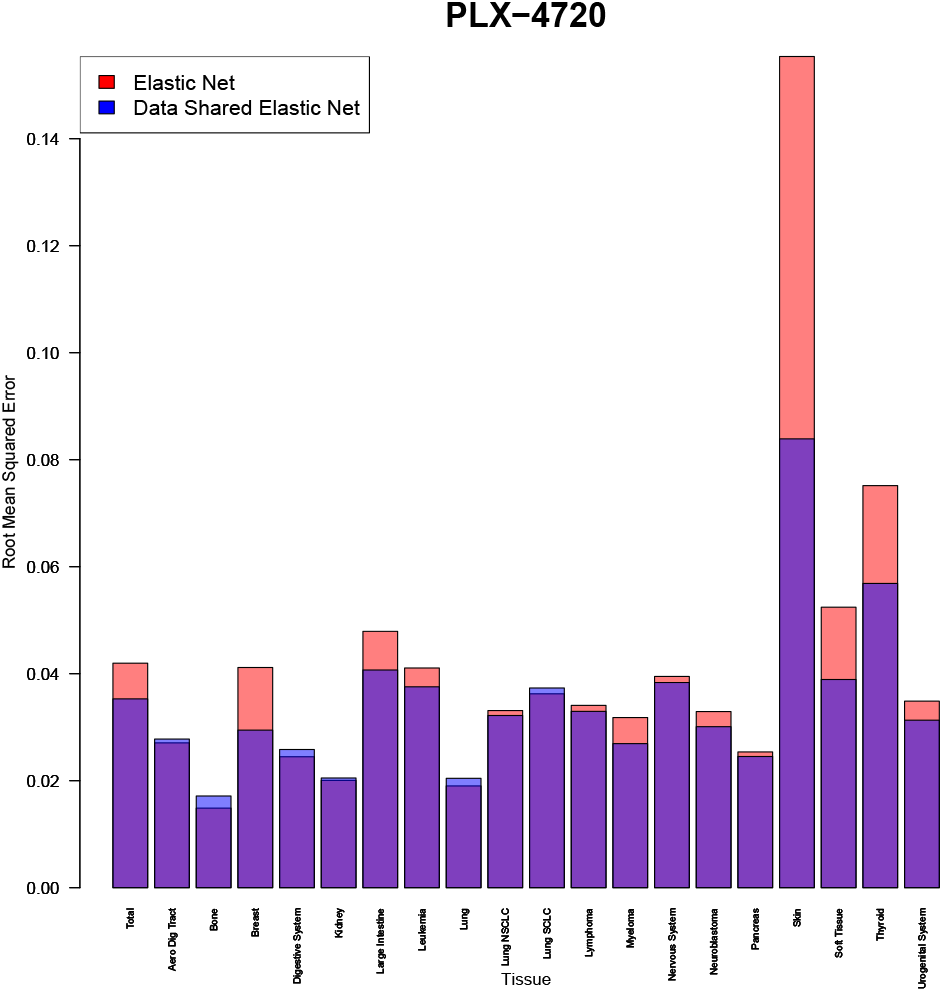
Comparison of RMSE between the DSEN and EN models for Predicted AUC values of PLX-4720. RMSE values were computed for all cell lines as well as within specific tissue types. The most substantial disparity in RMSE values is observed in cell lines derived from skin tissues. Previous research by [18] has demonstrated the sensitivity of melanoma to treatment with PLX-4720.

**Fig. 10.**
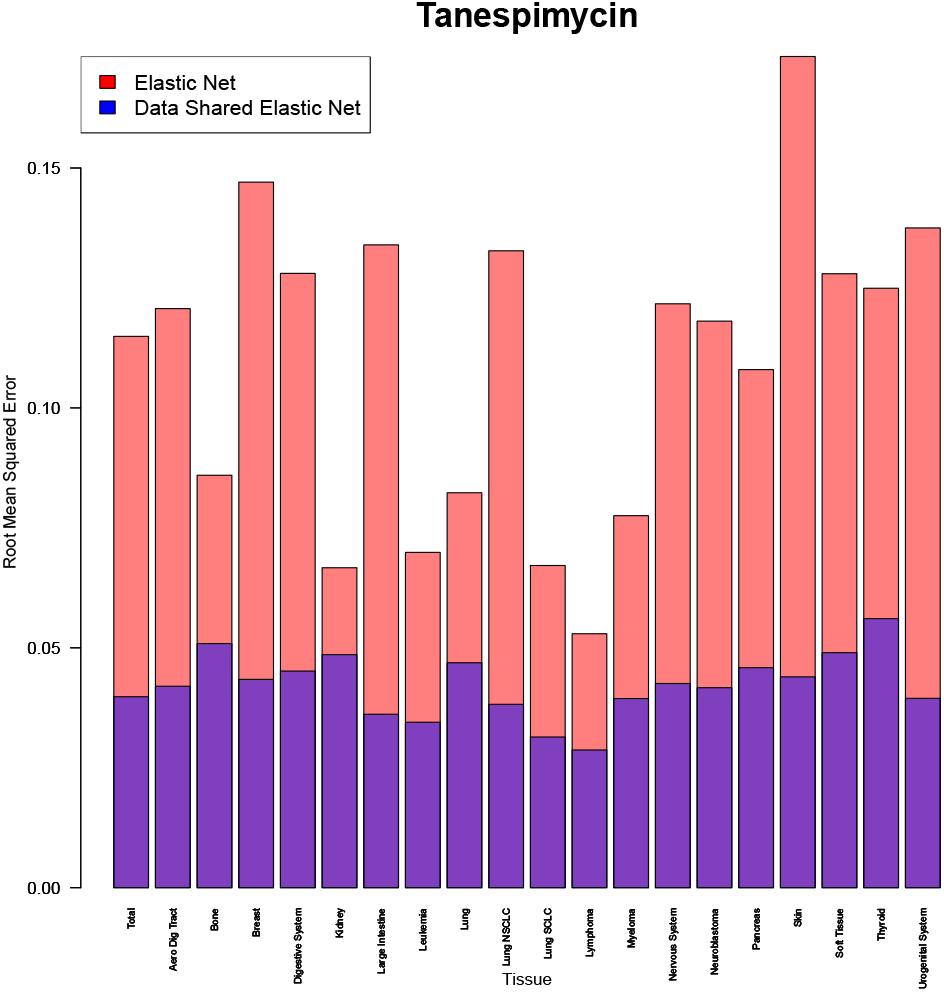
Comparison of RMSE Between the DSEN and EN models for Predicted AUC values of Tanespimycin. RMSE values were computed for all cell lines as well as within specific tissue types. The largest difference in RMSE is for cell lines derived from skin tissue. [14] showed that melanoma is sensitive to treatment with Tanespimycin.

To gain a more comprehensive understanding of the performance discrepancies between the DSEN and EN models, we created scatter plots of the observed AUC values in each cell line and their corresponding predicted values from both models. These visualizations, depicted in Figures 11, 12, and 13, allow for a clear assessment of the relationship between the sensitivity of cell lines and predicted AUC values.

**Fig. 11.**
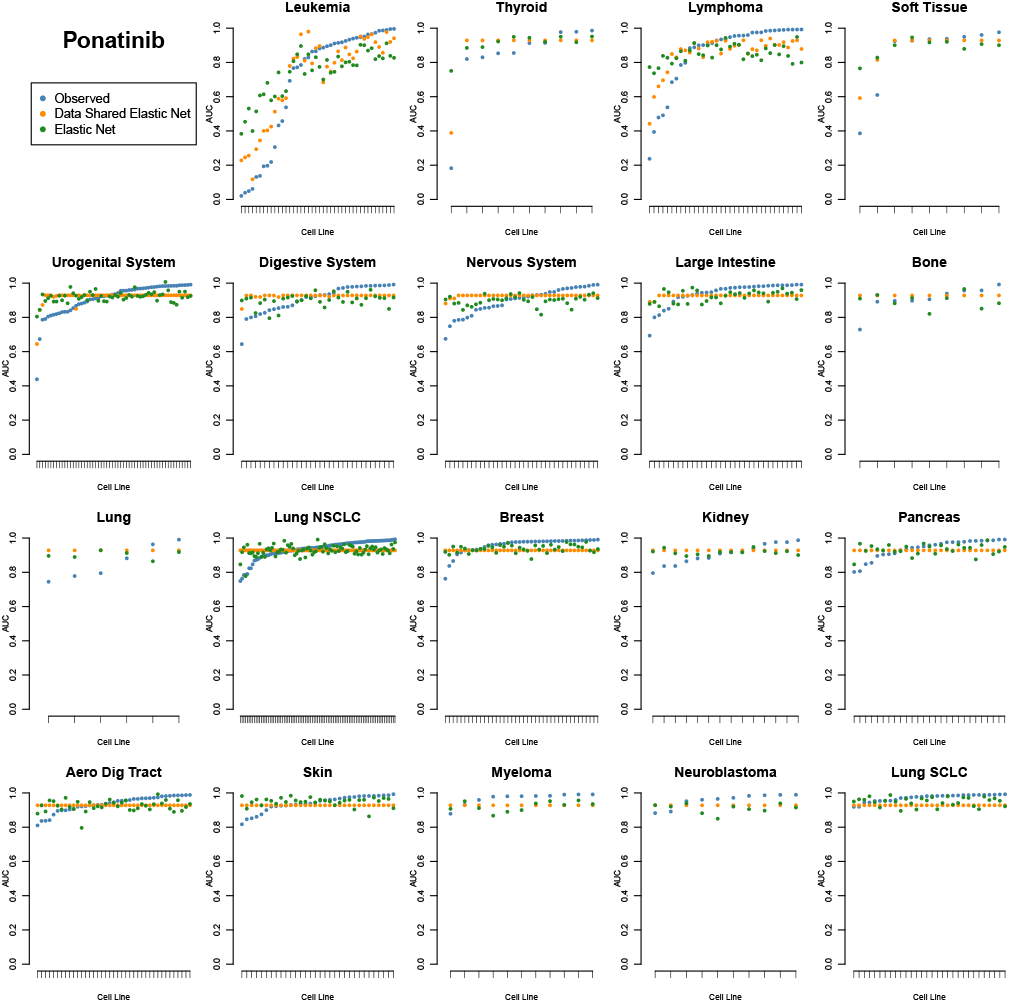
Comparison of Observed and Predicted AUC values per Cell Line for Ponatinib. Cell lines are arranged in ascending order based on their observed AUC values and grouped according to their tissue of origin. Notably, the EN model exhibits more conservative predictions compared to the DSEN model for cell lines displaying the highest sensitivity to Ponatinib treatment.

**Fig. 12.**
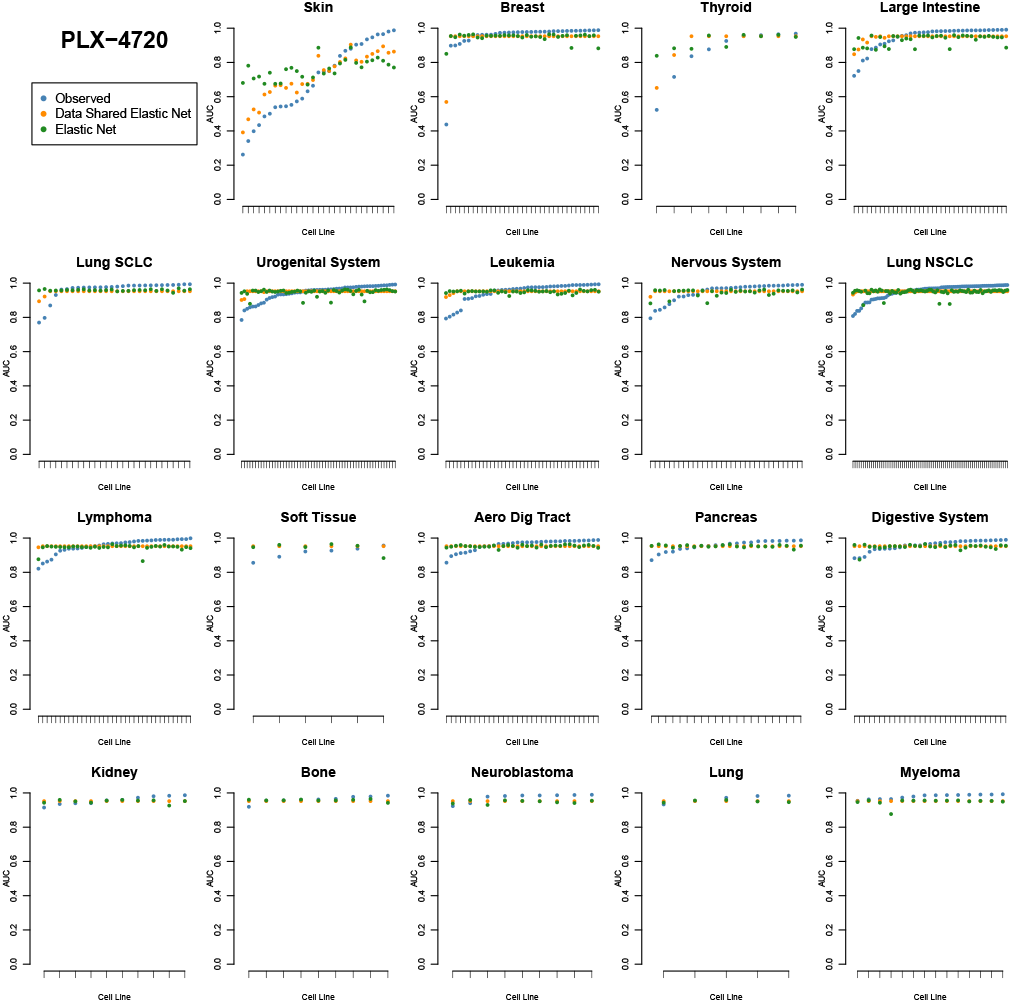
Comparison of Observed and Predicted AUC values per Cell Line for PLX-4720. Cell lines are arranged in ascending order based on their observed AUC values and grouped according to their tissue of origin. Notably, the EN model exhibits more conservative predictions compared to the DSEN model for cell lines displaying the highest sensitivity to PLX-4720 treatment.

**Fig. 13.**
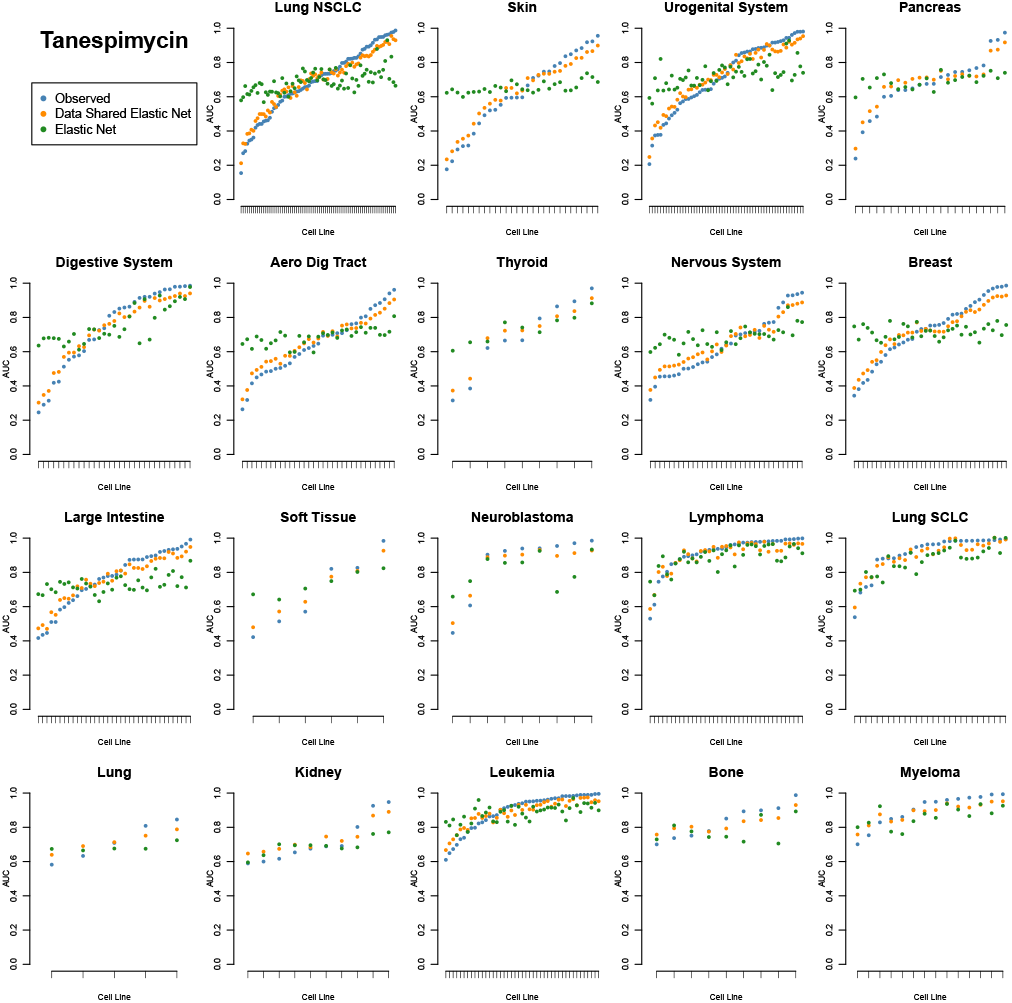
Comparison of Observed and Predicted AUC values per Cell Line for Tanespimycin. Cell lines are arranged in ascending order based on their observed AUC values and grouped according to their tissue of origin. Notably, the EN model exhibits more conservative predictions compared to the DSEN model for cell lines displaying the highest sensitivity to Tanespimycin treatment.

In the case of Ponatinib, cell lines derived from leukemic tissues exhibit the highest sensitivity. This is demonstrated by the lower AUC values in this specific tissue lineage. Importantly, the DSEN model’s predictions align more closely with the observed sensitivity to treatment compared to the EN model.

Similarly, for drugs PLX-4720 and Tanespimycin, it is observed that cell lines derived from skin tissues show the highest sensitivity. Once again, the DSEN model outperforms the EN model in accurately capturing and reflecting the observed sensitivity for cell lines that have the largest response to treatment.

To reinforce the finding that the most significant difference in performance between the DSEN and EN regressions lies in sensitive cell lines, we generated plots of the RMSE values for cell lines with an observed AUC less than or equal to a threshold value. These plots, illustrated in Figures 14, 15, and 16, demonstrate that when the threshold for cell line sensitivity is high, the DSEN model consistently outperforms the EN model. However, as the threshold for sensitivity is lowered, the performance of the EN model begins to resemble that of the DSEN model. This pattern holds for nearly all drugs that we have tested 17. These findings highlight the superior predictive performance of the DSEN model, particularly in capturing sensitivity patterns in cell lines that exhibit the largest response to treatment.

**Fig. 14.**
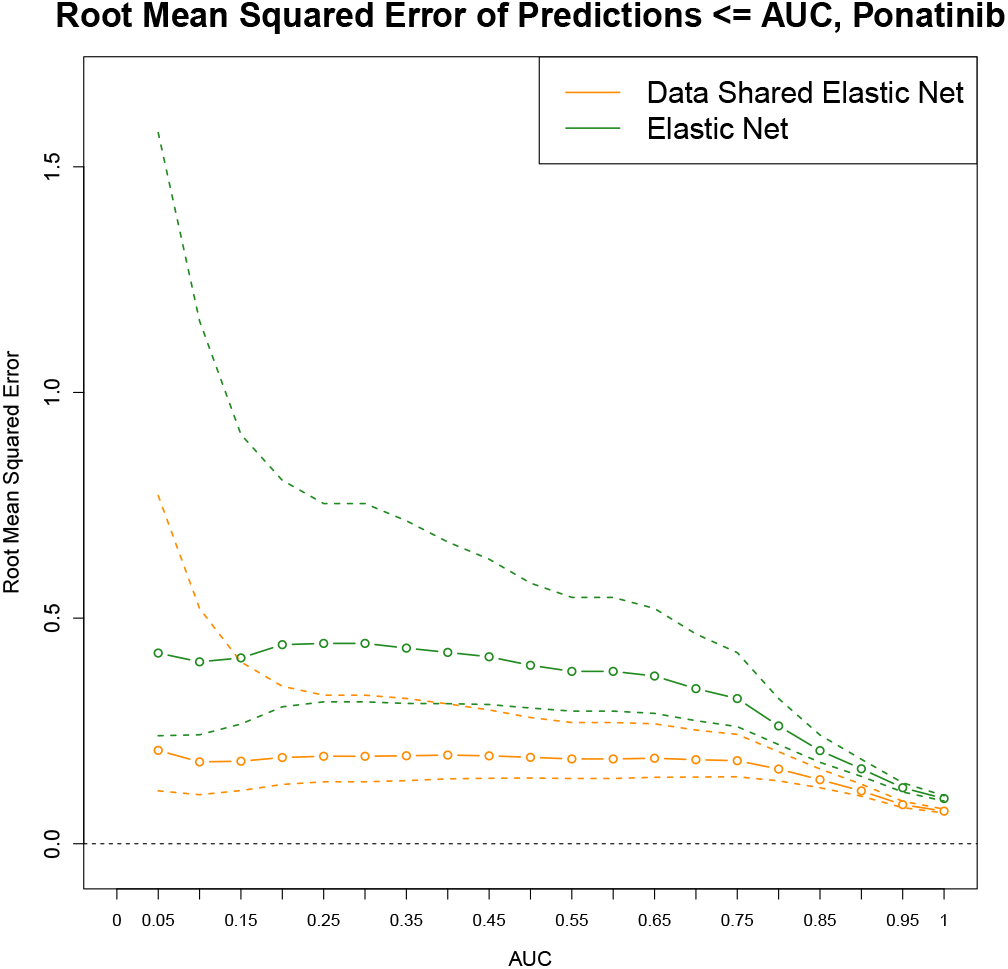
Comparison of RMSE between the DSEN and EN models for Ponatinib in cell lines with an observed AUC value less than or equal to a specified threshold. The dashed lines represent the 95% confidence intervals. The DSEN model demonstrates a lower RMSE than the EN model for cell lines exhibiting higher sensitivity to Ponatinib treatment.

**Fig. 15.**
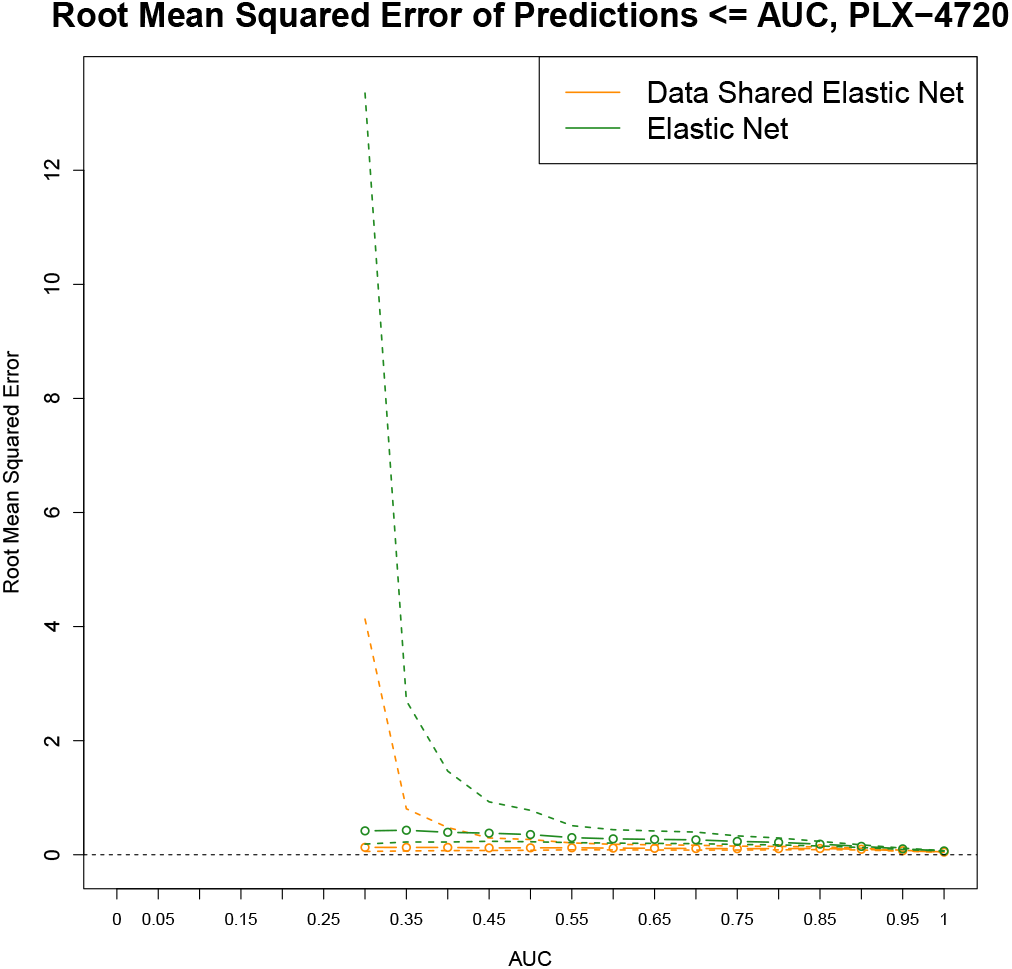
Comparison of RMSE between the DSEN and EN models for PLX-4720 in cell lines with an observed AUC value less than or equal to a specified threshold. The dashed lines represent the 95% confidence intervals. The DSEN model demonstrates a lower RMSE than the EN model for cell lines exhibiting higher sensitivity to PLX-4720 treatment.

**Fig. 16.**
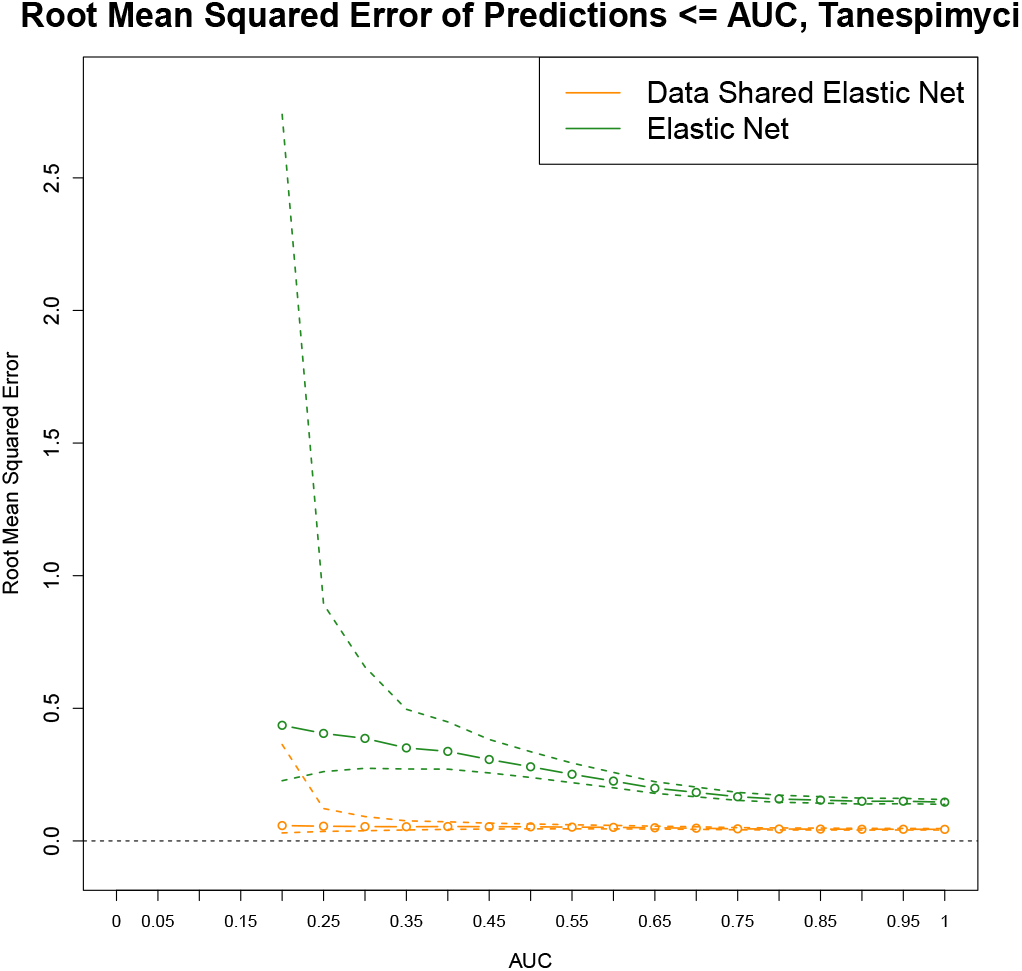
Comparison of RMSE between the DSEN and EN models for Tanespimycin in cell lines with an observed AUC value less than or equal to a specified threshold. The dashed lines represent the 95% confidence intervals. The DSEN model demonstrates a lower RMSE than the EN model for cell lines exhibiting higher sensitivity to Tanespimycin treatment.

### Inference from DSEN Regression

In both DSEN and EN regression, we can draw inferences regarding predictor importance using average coefficient estimates and the proportion of non-zero coefficient estimates from bootstrap re-samples. However, DSEN regression offers additional insights. It provides either a shared parameter estimate and the difference from the shared estimate for each tissue or tissue-specific coefficient estimates. For our chosen *r*_*t*_ value, a shared coefficient estimate exists only when all tissues have a non-zero coefficient estimate in the same direction for a particular feature. Otherwise, the shared coefficient estimate will be zero.

Figure 18 illustrates an example where only NQO1 expression has a shared parameter estimate, while other features do not. In our model, NQO1 expression is associated with increased sensitivity to treatment in all tissue types. This finding is supported by the fact that NQO1 expression increases binding to HSP90, the main target of Tanespimycin [10].

**Fig. 17.**
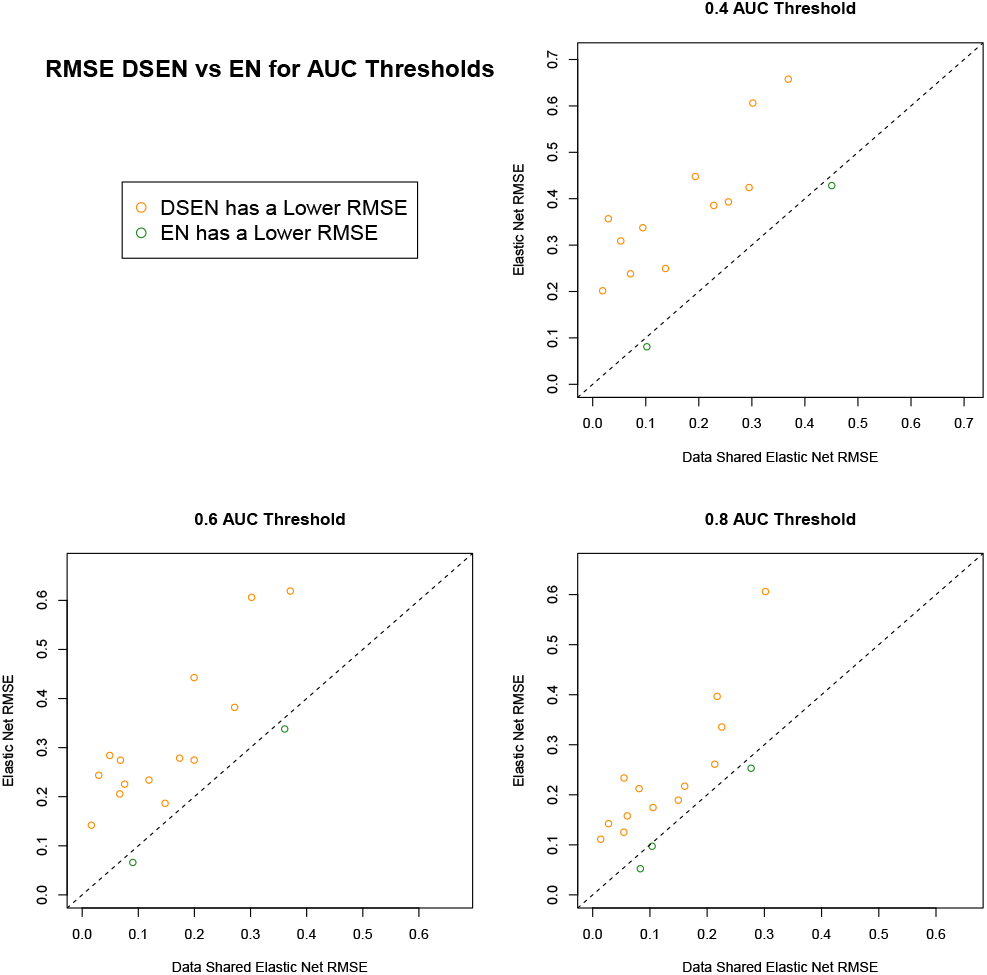
Comparing RMSE for Predicted Values for Cell Lines under an Observed AUC Threshold from the DSEN and EN models for All Drugs. A lower AUC value indicates higher drug sensitivity. Points above the y = x line indicate the DSEN model has a lower RMSE. Points below the y = x line indicate the Elastic Net model has a lower RMSE. The DSEN model outperforms the EN model, especially in cell lines that exhibit high sensitivity.

**Fig. 18.**
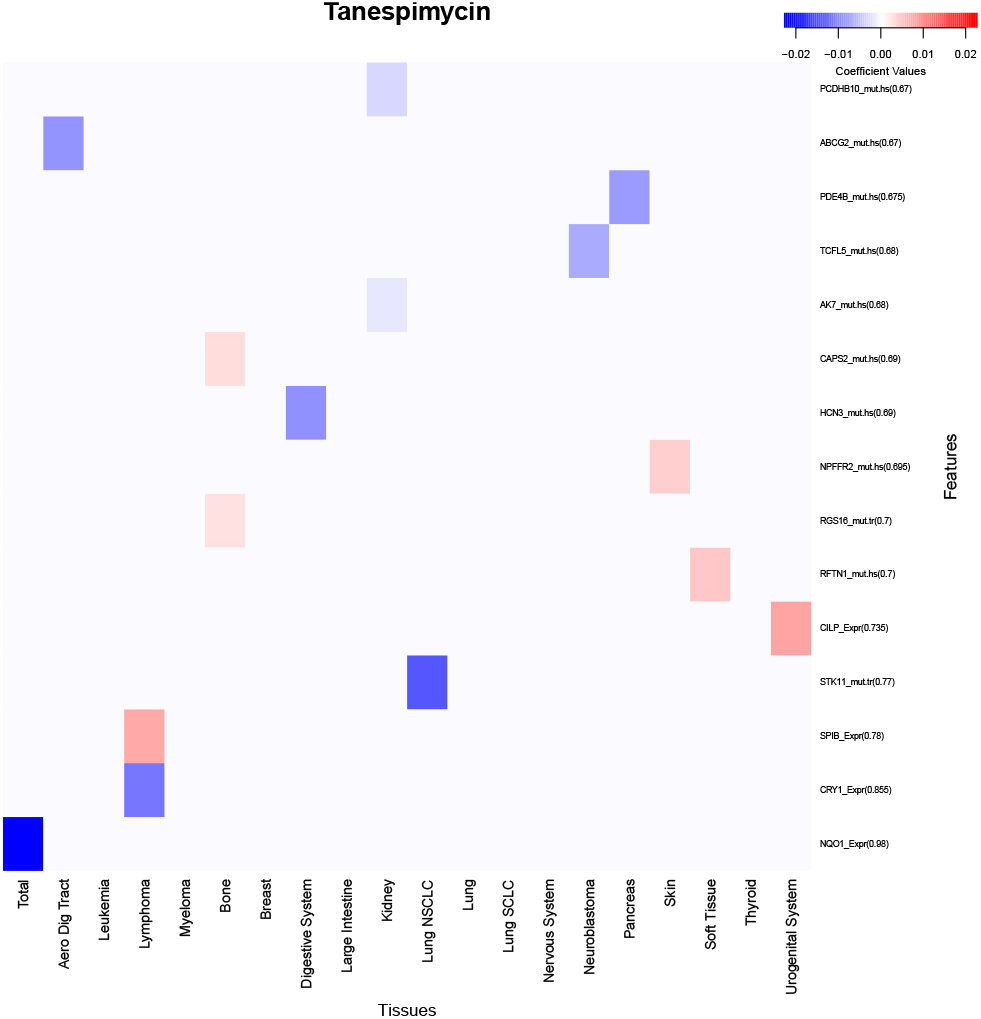
Heatmap of mean coefficient estimates from 200 bootstrap replicates of the DSEN model for Tanespimycin. The proportion of times the shared or tissue-specific coefficient estimates were non-zero in the bootstrap replicate is indicated in parenthesis beside the corresponding feature name. The 15 features with the highest non-zero proportion are presented. In our model, NQO1 expression is associated with increased sensitivity to Tanespimycin treatment in all tissue types.

The average coefficient estimate and the proportion of times the shared or tissue-specific coefficient estimate was non-zero were visualized in a heatmap for each drug. Due to space limitations, only the top 15 features with the highest non-zero proportion were included.

In Figure 19, SHP-2 protein expression is shown to be an important feature in predicting sensitivity to treatment with Ponatinib. Previous research has shown that GAB2, which activates SHP2, is crucial for leukemogenesis mediated by BCR-ABL1, the primary target of Ponatinib [9]. Similar results were obtained for Nilotinib and Saracatinib, which also target BCR-ABL1 [17] [6]. Furthermore, the top predictor for PLX-4720 sensitivity was identified as B-RAF mutation in Figure 20, aligning with the fact that B-Raf is the primary target of PLX-4720 [15].

**Fig. 19.**
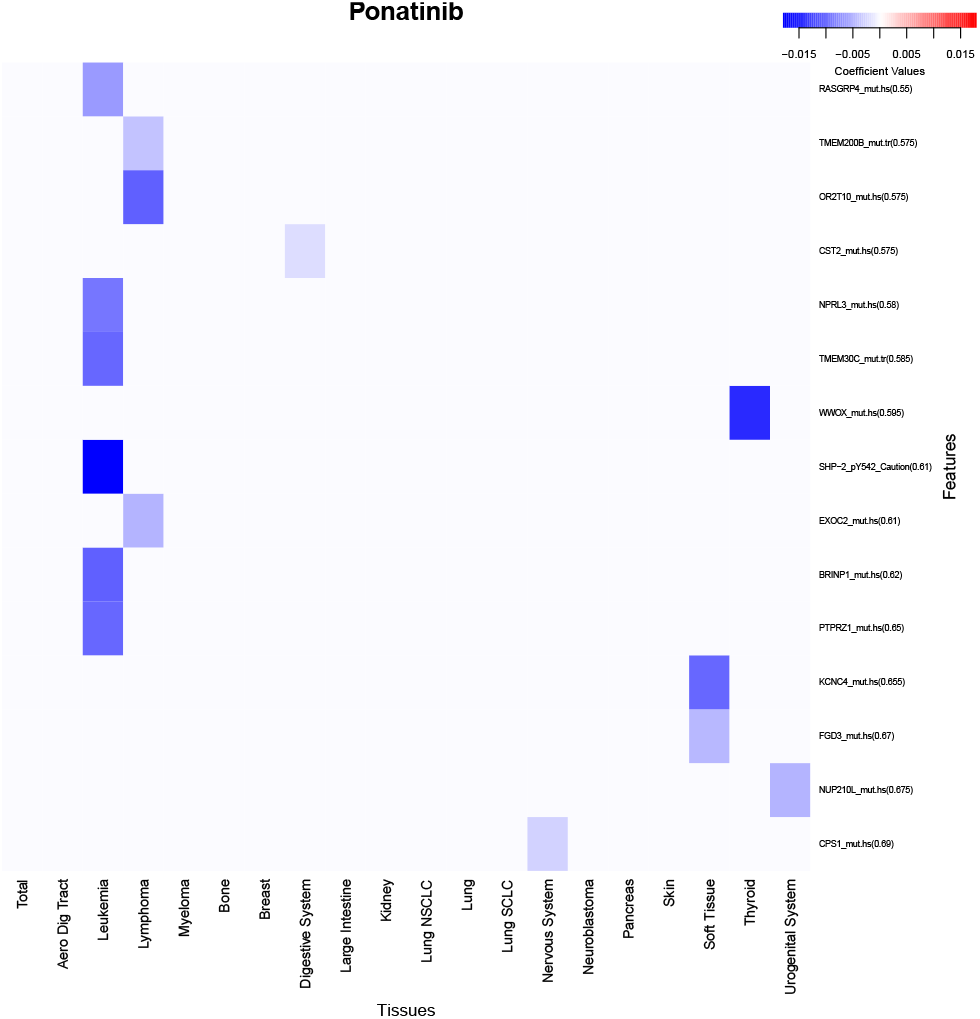
Heatmap of mean coefficient estimates from 200 bootstrap replicates of the DSEN model for Ponatinib. The proportion of times the shared or tissue-specific coefficient estimates were non-zero in the bootstrap replicate is indicated in parenthesis beside the corresponding feature name. The 15 features with the highest non-zero proportion are presented. SHP-2 expression has the largest negative mean coefficient estimate, indicating it is an important predictor for cell line sensitivity to Ponatinib. SHP-2 is necessary for leukemogenesis and is in the targeted pathway of Ponatinib [9].

**Fig. 20.**
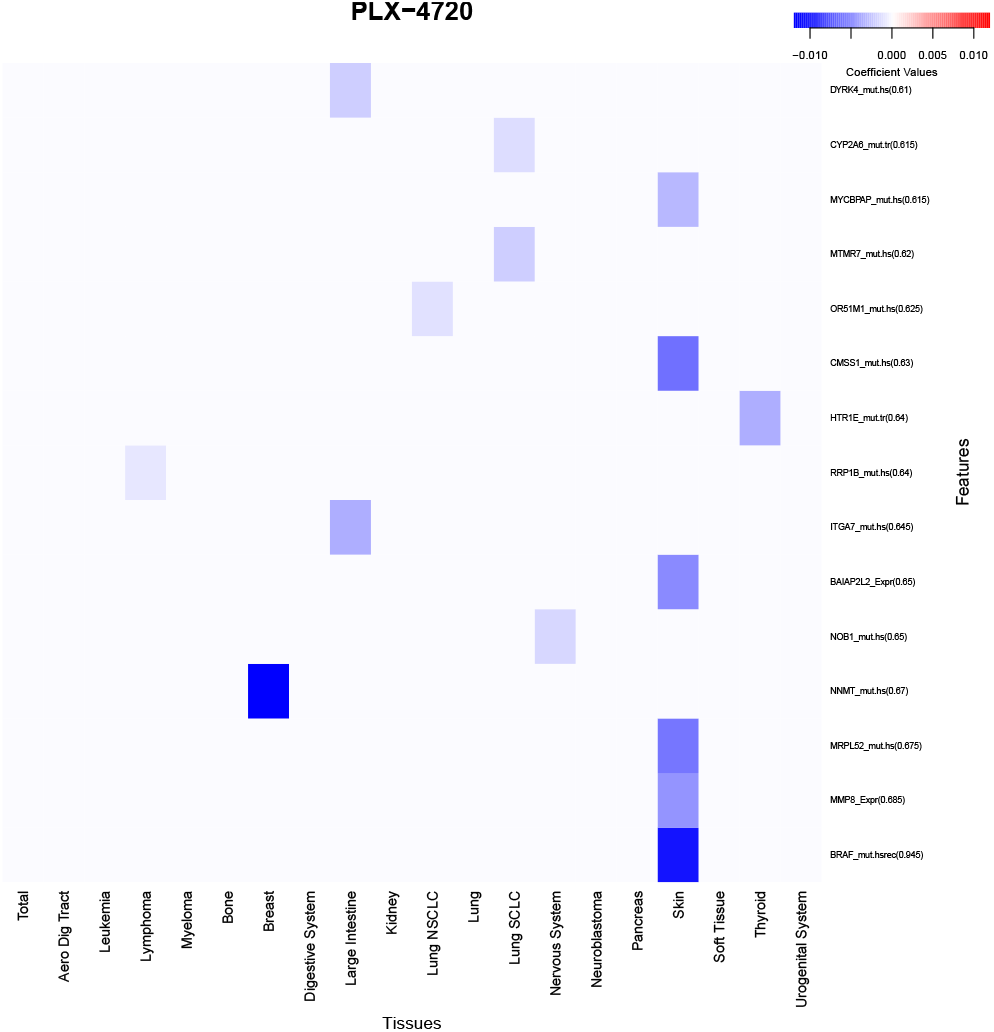
Heatmap of mean coefficient estimates from 200 bootstrap replicates of the DSEN model for PLX-4720. The proportion of times the shared or tissue-specific coefficient estimates were non-zero in the bootstrap replicate is indicated in parenthesis beside the corresponding feature name. The 15 features with the highest non-zero proportion are presented. B-Raf mutation expression has the largest negative mean coefficient estimate and proportion of non-zero coefficient estimates, indicating it is an important predictor for cell line sensitivity to PLX-4720. PLX-4720 has been shown to target thyroid cancer cells over-expressing B-Raf [15].

It is worth noting that EN regression also identified SHP-2 as a top predictor for Ponatinib sensitivity [7] and B-Raf as a top predictor for PLX-4720 sensitivity [3] in previous studies. These consistent findings across EN and DSEN regression demonstrate that both approaches are capable of yielding results that align with the underlying biological pathways of the treatments. However, DSEN regression provides an additional advantage by identifying the specific tissues where top predictors contribute the most valuable information regarding drug sensitivity.

## Discussion

The findings presented in the first portion of the results section shed light on the impact of data leakage on our models and the subsequent implications for drawing accurate conclusions. The overestimation of model stability due to data leakage becomes evident as the proportion of non-zero features in bootstrap re-samples tends to increase. Additionally, the magnitude of coefficient estimates will vary, although the sign of the estimate is generally preserved. This tendency towards overestimation can lead to unwarranted confidence in the importance of feature contributions to drug sensitivity. Furthermore, data leakage results in an overestimation of prediction accuracy during cross-validation and can contribute to overfitting by utilizing information from the test set during parameter tuning. Therefore, it is crucial to exercise caution and prevent data leakage to ensure the reliability and replicability of our models.

We showcased how a multitask learning framework can enhance prediction accuracy and inference when contrasted with single-task learning. Comparing Data Shared Elastic Net regression to Elastic Net regression, several advantages of the former become apparent. DSEN regression exhibits superior overall prediction accuracy, with the most substantial improvement observed for cell lines demonstrating high sensitivity to treatment. These advancements in accuracy bear significant implications for addressing scientific inquiries, such as predicting patient response to treatment or identifying potential drug candidates for cancer repurposing. False negatives, in particular, pose considerable concerns in these scenarios, as making overly conservative predictions can have severe consequences for patient care and hinder cost and time savings associated with drug repurposing. The adoption of DSEN regression helps mitigate these issues by minimizing false negatives, ultimately enhancing the quality of care and resource allocation.

Another notable benefit of utilizing DSEN regression lies in its ability to provide tissue-specific coefficient estimates. This feature facilitates the identification of cellular pathways that contribute significantly to drug sensitivity within distinct tissue types. Consequently, the model aids in discerning whether a drug operates through uniform pathways across all tissues or targets different pathways in each tissue, thereby advancing our understanding of drug mechanisms.

However, it is important to acknowledge the limitations of DSEN regression. A primary drawback is the requirement for all tasks to share the same set of features. Missingness within datasets further compounds the problem. In such cases, a feature missing in one dataset necessitates exclusion from all other datasets, even if the feature is available in those sets. Addressing missing data through imputation techniques can help alleviate this limitation. Additionally, the size of the feature matrix increases rapidly with the number of tasks, leading to storage difficulties and lengthy computation times. Employing sparse matrix representations and utilizing solvers designed to handle such matrices can offer a potential solution. Fortunately, the R package GLMNET provides the necessary functionality to address this challenge effectively.

A future direction for this project would be to evaluate model generalizability to other publicly available cancer cell line datasets. Model prediction accuracy is typically higher within the dataset the model was developed on than when the model is applied to another dataset. Applying our modeling procedure across datasets would allow us to ensure we are not overfitting our data and making over-optimistic estimates of the prediction accuracy. Moreover, an extension of our model to accommodate multi-output prediction would be valuable. Predicting multiple drug sensitivity measures simultaneously holds significant utility and can enhance the accuracy of predictions for each measure.

## Conclusion

In summary, Data Shared Elastic Net regression demonstrates superior prediction accuracy, mitigates false negatives, and facilitates tissue-specific analyses while maintaining the straightforward interpretability of Elastic Net regression. However, careful consideration must be given to the shared feature set requirement, as well as the challenges posed by missing data and computation efficiency. By navigating these limitations, researchers can harness the full potential of Data Shared Elastic Net regression to advance our understanding of drug sensitivity and enhance patient care.

